# Protease-driven remodeling of stabilization networks during adenovirus assembly

**DOI:** 10.64898/2026.08.04.740336

**Authors:** José Gallardo, Roberto Marabini, Marta Martínez, Carmen San Martín

## Abstract

At least thirteen different proteins form the adenovirus virion, including those bound to the dsDNA genome in the core. To produce infectious particles, the adenovirus protease (AVP) cleaves many of these proteins during genome packaging, using the viral genome as a cofactor. Here, we determine high-resolution structures of two types of adenovirus particles devoid of genome and core proteins and stalled at different AVP processing stages. We find that the N-terminal regions of penton base and internal minor coat protein IIIa are disordered when the core is absent and uncleaved packaging protein L1 52/55 kDa is present, likely enabling correction of early assembly errors at the vertex. Assignment of previously unmodeled densities reveals that proteins IIIa and VIII form long-range bridges linking vertex capsomers to the facet center before proteolytic maturation. Cleavage by AVP remodels these connections, diminishing capsid-wide stabilizing interactions and promoting the metastable state that prepares the mature virion for uncoating.

**Significance statement:** Adenoviruses cause disease and are major platforms for gene delivery, yet how their capsids are built and primed for infection remains incompletely understood. High-resolution cryo-EM structures of empty particles arrested at successive proteolytic maturation stages show previously unrecognized interactions that regulate capsid stability. Uncleaved packaging protein L1 52/55 kDa induces disorder at the icosahedral vertex during early assembly, likely facilitating error correction, while minor coat proteins IIIa and VIII serve as transient scaffolds stabilizing the capsid before proteolytic remodeling. Maturation thereby converts a stable assembly intermediate into a metastable particle ready for uncoating. These findings reveal how structural plasticity of minor coat proteins coordinates adenovirus assembly, maturation, and infectivity.

## Introduction

Adenoviruses assemble non-enveloped, *pseudo*-T=25 icosahedral particles containing a linear, double stranded DNA genome (∼35 kbp in human adenoviruses, HAdVs). The virion is composed of the major coat protein (hexon); penton base and fiber at the vertices; internal minor coat proteins (IIIa, VI, VIII); and DNA-condensing proteins in the core (VII, µ) (1). Mastadenoviruses (including HAdVs) also incorporate genus-specific external minor coat protein IX and core protein V (2). The icosahedral shell is formed by two types of tiles: a *Group of Nine* hexon trimers (GON) forming the central plate of facet, and a *Group of Six* capsomers (GOS) formed by each penton and its peripentonal hexons (3). Infectious particles also contain genome packaging/replication proteins (IVa2, terminal protein TP), and the DNA-binding adenovirus maturation protease (AVP) (4). Other assembly factors (proteins L1 52/55 kDa, L4 100 kDa, and L4 33/22 kDa) are absent or barely detectable in mature particles (5).

Packaging protein L1 52/55 kDa interacts with capsid, core, and other packaging proteins, and also undergoes self-association, serving as a molecular tether between capsid and core (6–8). During maturation, AVP cleaves capsid (IIIa, VI, VIII), core (VII, µ, TP), and L1 52/55 kDa proteins (7, 9). As a result, the capsid-core linkage established by L1 52/55 kDa is removed, internal pressure increases, and the vertex region is destabilized. The particle thereby becomes metastable and primed for sequential uncoating, with penton release representing the earliest detectable structural change (10, 11). Maturation likely coincides with packaging, as DNA activates AVP (4). The human adenovirus type 2 (HAdV-C2) *ts1* mutant produces fully packaged particles where no maturation cleavages have occurred, and serves as a model for structure and function of immature adenovirus particles (11–13).

Adenovirus naturally forms empty capsids (14), also referred to as light particles, as they have lower density than their full counterparts (heavy particles). Light particles may represent assembly failures rather than intermediates (15). The concerted adenovirus assembly model (16, 17) proposes that protein L1 52/55 kDa recruits capsid fragments around the genome condensed by core proteins (**Figure 1.I**). As the capsid grows, cleavage by AVP begins (**Figure 1.II**). If the genome-capsid tether fails, empty particles lacking the genome and core proteins form (**Figure 1.III-IV**), containing AVP targets at varying cleavage states, among them L1 52/55 kDa. Here we refer to empty (E) capsids as E0 when AVP cleavages are not detected, and E1 when they are. Assembly failure in the *ts1* mutant necessarily produces E0 capsids, as the protease is not encapsidated (18). The packaging-delayed mutant HAdV-C5 FC31 produces E1 particles with partially processed AVP targets (15, 19). HAdV-C2 *ts1* also produces full (F) immature capsids (F0, particle with genome and core proteins but no cleavages) (**Figure 1.V**). Mature virions (F1, fully packaged and cleaved) assemble only when core-capsid interactions are appropriately coordinated (**Figure 1.VI**) (16).

**Figure 1.**
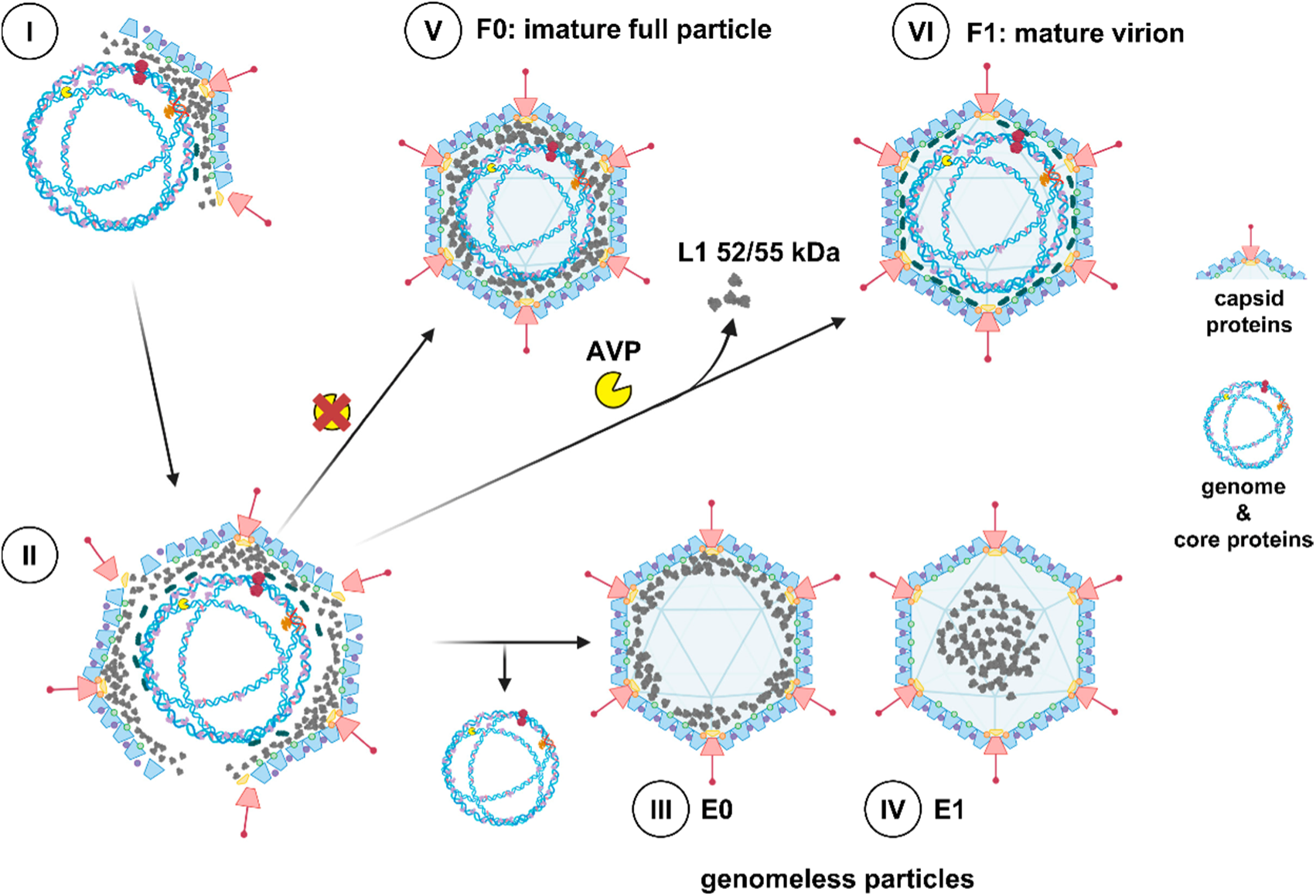
Schematic representation of the adenovirus concerted assembly and packaging model. (**I and II**) Capsid recruitment to the core mediated by L1 52/55 kDa protein. (**III and IV**) Failed assembly products containing unprocessed (E0) or partially processed (E1) forms of protein L1 52/55 kDa, but lacking genome and core proteins. (V) Genome-containing, immature particles (F0) resulting from lack of AVP activity. (VI) Mature particle (F1), assembled when both genome incorporation and AVP processing occur correctly. Created in BioRender. https://BioRender.com/ry7dtbp

High resolution structures are available for the mature HAdV-C5 virion (20) and for immature, fully packaged particles produced by a HAdV-C5 mutant (Ad5F35-ts1) containing the *ts1* mutation (21). The latter showed large conformational differences between the mature and immature versions of protein VIII, and an additional ordered appendage domain (APD) in protein IIIa. To provide information on the architecture of earlier assembly intermediates, here we report high-resolution structures of HAdV-C2/5 E0 and E1 particles, and compare them with their genome-containing counterparts. We observe changes in proteins IIIa and penton base related to the presence of the core and/or packaging protein L1 52/55 kDa. We also locate previously unassigned fragments of proteins VIII and IIIa, showing how these proteins contribute to vertex preparation for uncoating upon maturation. These results provide new details on the changes undergone by adenovirus capsid components during morphogenesis, and how they relate with stability modulation of the viral particle.

## Results

### High-resolution structure of E0 and E1 particles

Details on the specific specimens used in this work for E0 and E1 capsids, as well as the previously reported F0 and F1 structures used for comparison, are shown in **Table 1**. Denaturing electrophoresis followed by silver staining showed that both E0 and E1 particles lack core proteins V and VII (**Figure S1A**), as expected for genomeless particles. In agreement with impaired proteolytic processing, E0 particles presented the precursor forms of proteins VIII (pVIII) and VI (pVI), while E1 contained an intermediate form of VI (iVI, cleaved at one of its two AVP consensus sites), and mature VI (9, 15) (**Figure S1B**). E0 contained full-length protein L1 52/55 kDa, while E1 particles showed a small amount of unprocessed L1 52/55 kDa and several populations of cleaved products (**Figure S1**) (7, 15).

**Table 1.**
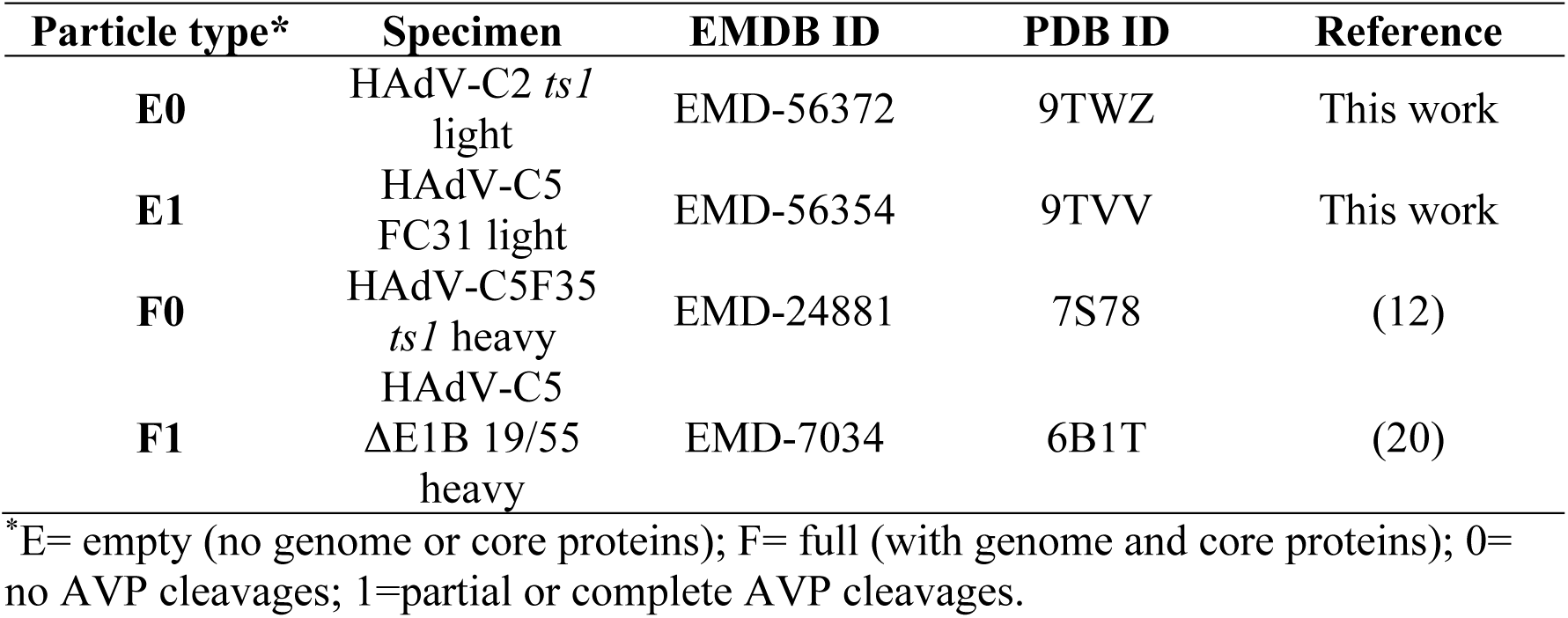
Specimens compared in this work.

High-resolution cryo-EM maps were obtained for E0 (3.3 Å) and E1 (3.0 Å) particles (**Figure S2, Table S1**). Molecular models for hexon, penton base, proteins IIIa, and VIII were traced, along with some copies of the protein VI N-terminal peptide pVI_N_ (20) (**Figure 2, Tables S2 and S3**). Since the density for polypeptide IX was weak in E1 particles, this protein was traced only in the E0 map. Some additional small densities were initially unassigned but were of sufficient quality to be interpreted later on (see below) (**Figure 2, unassigned**). Radial average profiles of the maps showed low signal for capsid contents, consistent with the lack of genome and core proteins (**Figure S3**). As the specimen used for E0 capsids was HAdV-C2 *ts1* light particles (**Table 1**), E0 is the first reported high-resolution structure of the HAdV-C2 icosahedral shell. HAdV-C5 and –C2 have a genome identity of 98.68% with a 97% query cover (BLAST). Sequence differences in the capsid proteins were considered during model building.

**Figure 2.**
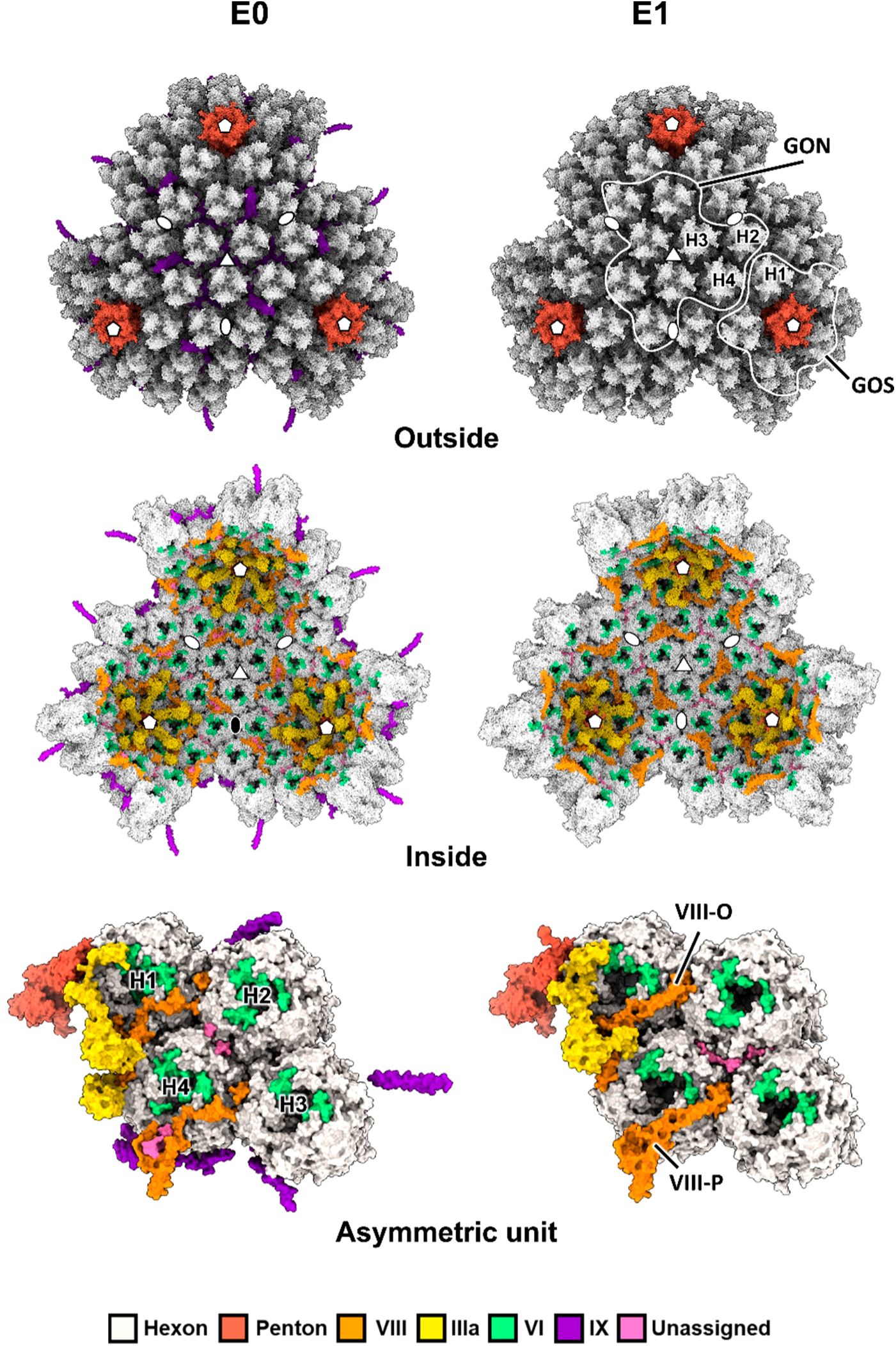
Structure of adenovirus E0 and E1 particles. Traced proteins in one facet and neighboring asymmetric units (AU) are shown from outside and from inside the particle, with a zoom into the interior of one AU on the bottom. Hexon trimers in one AU are labeled H1 to H4. Pentagons, triangles, and ovals indicate the icosahedral 5-fold, 3-fold, and 2-fold symmetry axes, respectively. One GOS and one GON are delimited with white shapes. Models traced in initially unassigned densities (two in E0, one in E1) and the positions of the two independent copies of protein VIII (chains O and P) are also shown.

### Hexon, penton base and protein IX

Hexon and penton base structures in E0 and E1 particles were very similar to their previously reported genome-containing counterparts, F0 and F1 (**Figure 3A and S4**). Differences found when comparing E0 and F0 hexons were concentrated in the loops exposed on the capsid surface (**Figure 3A**), and can be attributed to differences in sequence between HAdV-C5 and –C2 rather than packaging state, since hexons present the lowest identity (87%) of all structural proteins composing the HAdV-C2 and HAdV-C5 viral particles. These results indicate that the hexon structure is not affected by genome presence.

**Figure 3.**
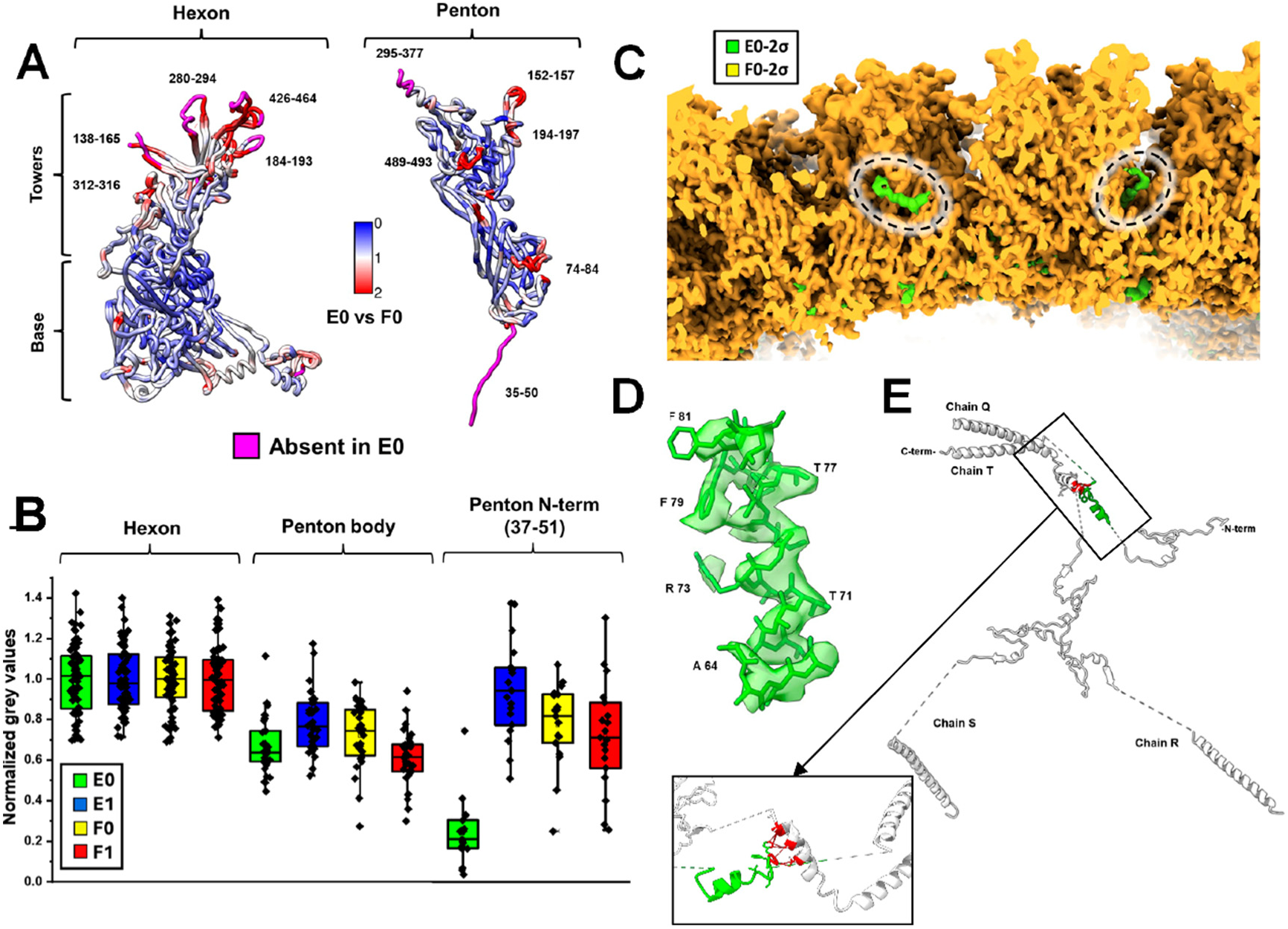
Structure of major capsid components and external cementing protein **IX**. (**A**) Comparison between hexon and penton base monomers in E0 and F0 particles, colored by RMSD (scale bar in Å). Residue numbers indicate the regions with the largest differences between specimens. Ordered residues traced in F0 but absent in E0 particles are depicted in pink. **(B)** Quantitative comparison of grey levels in map regions corresponding to hexon and penton base proteins. Each dot corresponds to the map density value at the position of a marker in UCSF Chimera, normalized to the hexon density. **(C)** Subtraction of F0 from E0 unsharpened maps showing extra density between hexons on the outer capsid surface (dashed ovals). Both maps are rendered at 2σ threshold above the average density. **(D)** Protein IX residues 64-82 fit into the extra density. **(E)** The four IX copies present in the asymmetric unit are depicted in white, with the newly traced helix in green and its contacting residues in a neighboring rope domain in red (bottom right, slightly rotated view).

For penton base, the main difference among the four specimens compared was the loss of order in its N-terminal region (residues 37-51), which could not be traced in the E0 map (**Figure 3A**). To quantify this difference, we compared grey levels (density values) between the four maps. When grey levels for hexon were considered the standard (density level = 1), the main body of penton base (comprising the single jelly roll domain) yielded values near 0.7. This value was very similar for the four compared maps, regardless of their packaging or maturation state. However, when only the map region corresponding to the N-terminal tail of penton base was considered, there was a large decrease in density only in the E0 particles, with grey values dropping to 0.2 (**Figure 3B**). The mean and standard deviation of density values in the region of interest for each map were used to test the null hypothesis that the four datasets have the same density. An ANOVA test yielded a p-value of 1.2 × 10⁻^14^ for the N-terminal region, indicating that at least one dataset differs significantly from the others. To determine which dataset(s) were different, a Tukey mean difference test was used, where all pairs of datasets were compared (**Table S4**). This analysis showed that at a significance level set to 0.05, grey values for the penton base N-terminal region in E0 differ clearly from all the other datasets. This result indicates that the penton base N-terminal region requires at least one of the following to become ordered: core presence, or AVP activity. Disorder in this region is observed only when neither genome packaging nor AVP target processing has taken place.

Protein IX is the only outer cementing protein in HAdVs, and has a complex oligomerization pattern to form a stabilizing network around hexons. N-terminal domains of three IX copies form triskelion-like structures, while the C-terminal domains of four IX copies form a coiled coil with three parallel helices and one antiparallel helix (3). The connections between triskelions and coiled coils, denoted as rope domains (residues 57-100), are flexible and difficult to trace confidently. In previous structures, only one or two copies out of the four rope domains present in the asymmetric unit have been modeled (12, 20). In our E0 map, we traced one helix in the rope domain of a protein IX molecule for which the rope domain had not been traced previously (chain T) (**Figure 3C-D, Table S3**). The newly traced helix does not appear in F0 particles, indicating that its ordering is not specific to immature particles. It is also unlikely that it is caused by a difference between HAdV-C2 and 5, since the protein sequence for the two viruses in this region and in the surrounding hexons is identical. This helix in the rope domain of protein IX chain T (residues 79-81) is close enough to the rope domain of chain Q (residues 68-76) to establish direct interactions between the two molecules (**Figure 3E**). This observation adds more interactions to those previously described for the well-ordered regions of the IX network: apart from the trimeric interaction present in the triskelion and the 4-chain interaction in the helix bundle, chains Q and T can dimerize through the rope domain.

### Identification of the central peptide of protein VIII

There are two copies of protein VIII on the internal surface of each asymmetric unit: one lining the GOS (chain O, at the boundary between hexons 1, 2 and 4), and a second one beneath the GON hexons (chain P) (**Figure 2**). Maturation of VIII produces two large fragments: N-terminal (Met1-Gly111) and C-terminal (Gly158-Asp227). The central peptide (residues 112 to 157) is further cleaved at Arg131 (9), with evidence that at least the 132-157 peptide remains within the viral particle (22). The previously reported F0 structure (12) revealed that maturation causes a large conformational change in protein VIII. We observe the same change in our E0 and E1 structures (**Figure 4A**). The E0 particle presented VIII in its immature form, while in E1 particles, VIII presented its mature form, with few variations from their respective genome-containing counterparts. In E0, although both parts of protein VIII are connected because no proteolytic cleavage has occurred, a large region in the N-terminal fragment (from residue 64 onwards) appears disordered. Upon cleavage by AVP (E1 particle), residues 64 to 111 become ordered, and the tracing is instead interrupted at the cleavage sites (residues 111 and 158), consistent with the separation of the excised central peptides (3, 20). When the N-terminal fragment becomes ordered, a small β-sheet is formed in the so-called neck domain, which was instead a helix in the immature form of protein VIII. This result is consistent with the biochemical analyses showing that in E1 particles, protein VIII has undergone cleavage by AVP (**Figure S1**) (15). It also indicates that the conformational change is uniquely caused by proteolytic processing and has no relation to the presence or absence of the genome.

**Figure 4.**
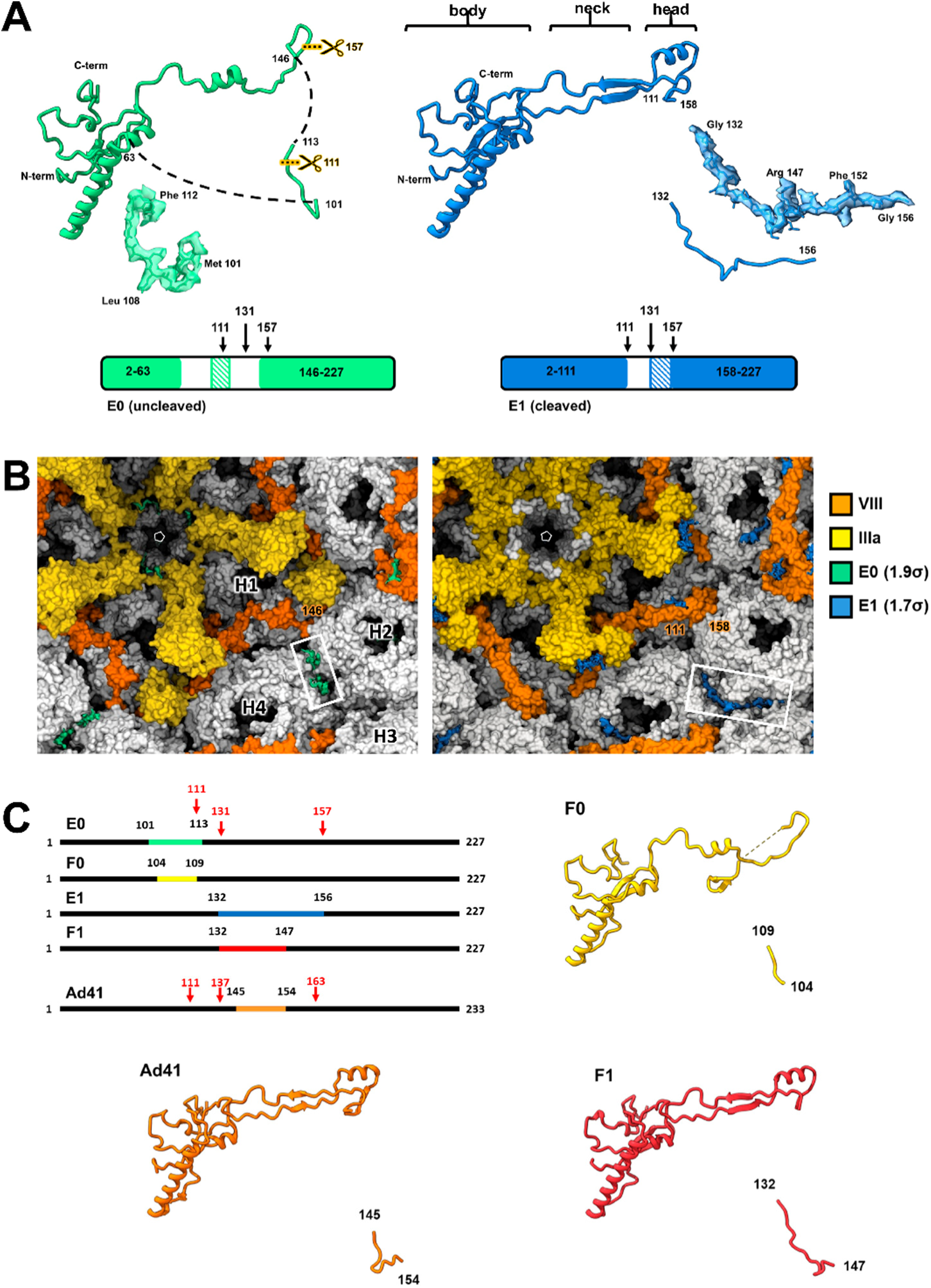
Protein VIII in E0 and E1 particles. **(A)** Structures of protein VIII in E0 and E1, showing the conformational change induced by AVP cleavage. The body, neck, and head domains are indicated, and the regions traced in each case are shown at the bottom, with arrows marking AVP processing sites, and diagonally hatched boxes indicating the peptides assigned using ModelAngelo. The fit of these peptides into the density is also shown. **(B)** Initially unassigned densities (unsharpened maps) near protein VIII (white frames). Residue labels indicate the position of the first traced residue of the protein C-terminal fragment in E0 (Leu146), and the position of the AVP processing sites (residues 111 and 158) in E1. **(C)** ModelAngelo identification of the VIII central peptide in adenovirus cryo-EM maps. Bars represent the protein sequences, highlighting the regions identified and the residues where AVP cleavages occur (red). Previously reported structures (F0, F1, Ad41) are shown, together with the residues identified in the central peptide.

In cryo-EM maps of adenovirus, relatively strong but small density features are often observed that cannot be clearly attributed to any traced protein. These densities are typically left unassigned or assigned only tentatively. In E0 particles, one such extra density is located between hexons H2 and H4. In E1 particles, the equivalent position is occupied by a longer, L-shaped density that extends along the H2-H3 interface (**Figure 4B**). When comparing with genome-containing maps, we observed that F0 presented the same extra density as E0, and F1 presented the same density as E1, suggesting that these densities correspond to a protein undergoing AVP processing. A similar L-shaped density has been reported in previous adenovirus structures and left unassigned in most of them, except in HAdV-F41, where it was attributed to core protein V (23). However, since there are no core proteins in our E0 and E1 particles (**Figure S1**), we interpret that a protein other than V is filling this position.

The extra density was located near the AVP cleavage sites of protein VIII (residues 111 and 158, chain O, under the vertex) in E1 particles, and even closer to the first traced residue in the C-terminal half of unprocessed protein VIII (residue 146) in E0 particles (**Figure 4B**). Running ModelAngelo (24) on E0, E1, F0, and F1 maps without providing an initial sequence yielded a match with protein VIII residues 103-111 in E0. When we applied the same procedure to a mature HAdV-F41 map (25), ModelAngelo identified residues 145-151 corresponding to the central peptide of VIII in the enteric virus. When provided with the protein VIII sequence, ModelAngelo assigned the following residues: 101-113 in E0, 104-109 in F0, 132-156 in E1, 132-147 in F1, and 145-154 in HAdV-F41 (**Figure 4C**). These results show that two different regions of protein VIII occupy the previously unassigned densities: residues 101-113 when AVP processing has not occurred (E0 and F0), and residues 132-156 in all the rest (or the equivalent region in HAdV-F41). In immature VIII, the identified region precedes and contains the first cleavage target by AVP, whereas in the particles where VIII has been cleaved, this region is displaced by the second peptide excised from the central part of the sequence.

Further support for the assignment of the central peptide comes from the observation that, in the E0 map, weak density connects the main body of unprocessed protein VIII (chain O) with the central peptide, possibly arising from the disordered region in the N-terminal fragment (Thr64-Gly111) (**Figure S5A, white rectangle**). No such connecting density was observed in the E1 map, even at very low rendering threshold. This observation is consistent with the fact that after cleavage, the central peptide is disconnected from the main body of the protein, and the disordered region of the N-terminal fragment generating the weak density in E0 has become ordered. Features similar to those described above for the peripentonal copy of VIII are also observed for the second independent copy of this protein (chain P, under the GON), with the density for the central peptide in E1 corresponding to the average of three equivalent chains located at the icosahedral 3-fold axis (**Figure S5B**). These results indicate that AVP processing, apart from driving a conformational change in the two large fragments of protein VIII, also reaccommodates the central peptide between hexons, and disrupts a link between the GOS and the hexons in the central plate of the facet (H2 and H4).

### Varying degrees of order in protein IIIa

There are five copies of protein IIIa beneath each vertex, gluing together penton and peripentonal hexons (**Figure 2**). Previous reports defined five structural domains in IIIa: GOS-glue domain, connecting-helix, VIII-binding domain, core-proximal domain (3), and APD (12) (**Figure 5A**). The GOS-glue domain starts at the N-terminus and is located under the penton, followed by the connecting helix and the VIII-binding domain, which is positioned under the VIII copy closer to the penton (chain O). The core-proximal domain is the last region traced in most adenovirus structures, and forms a long helix-turn-short helix structure (Asp251-Leu300) oriented towards the core (20). Downstream of the core-proximal domain, the APD has been observed only in F0 for HAdV-C5 (12), and in mature HAdV-D26 (26).

**Figure 5.**
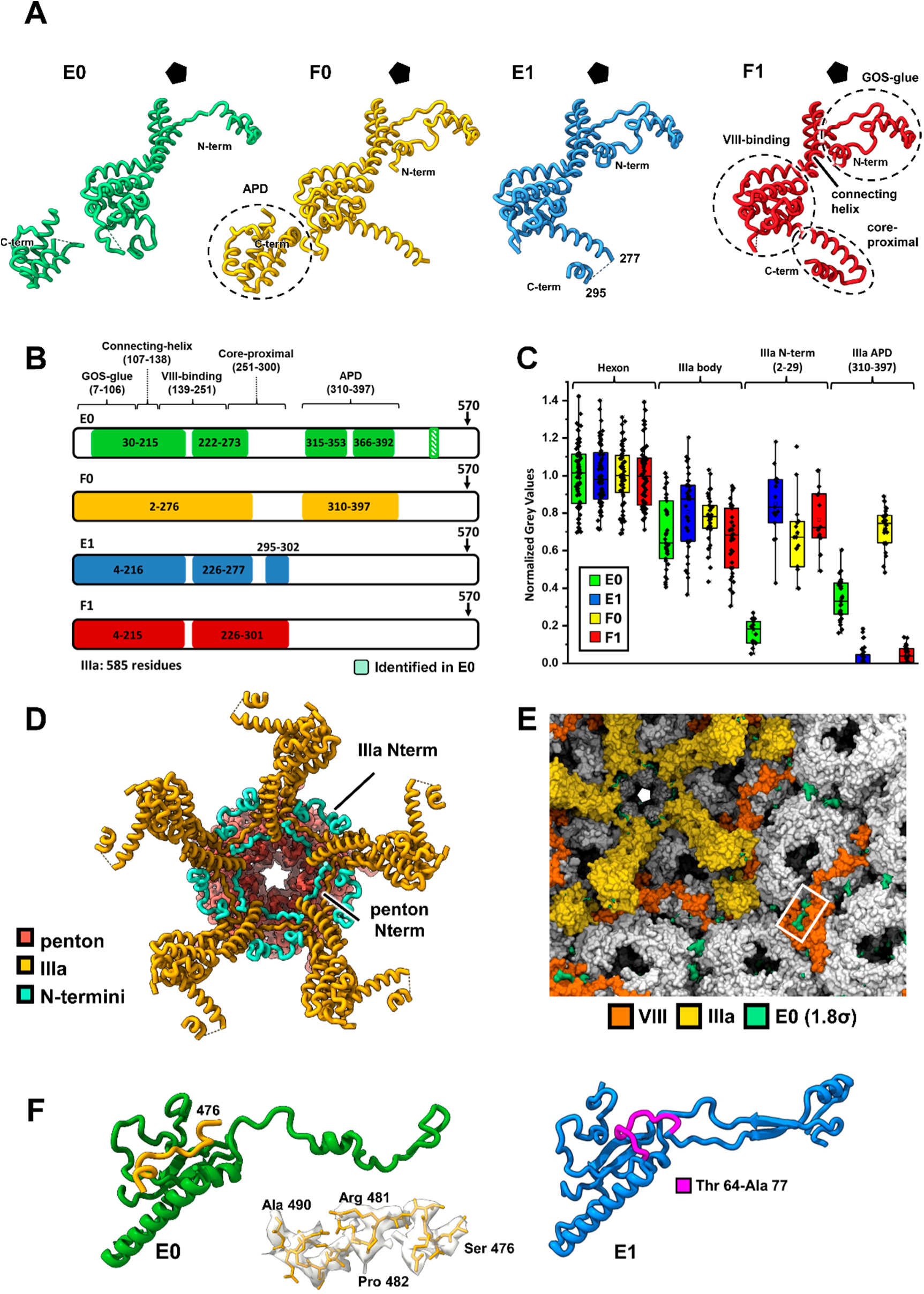
Structures of protein IIIa. **(A)** Molecular models traced in each of the four maps compared. **(B)** Regions of IIIa traced in each map. Arrows mark the AVP processing site. IIIa domains are also indicated in (A) and (B) as previously described (3, 12). The position of the peptide assigned using ModelAngelo is indicated with a diagonally hatched box. **(C)** Comparison of map grey level values for IIIa, normalized to those of hexon. Values for densities of the IIIa body were taken from the VIII-binding domain, connecting-helix domain, and final residues of the GOS-glue domain. **(D)** Penton base and IIIa proteins traced in E1 viewed from inside the capsid, with the residues disordered in E0 colored in aquamarine, to highlight their proximity in the context of the viral particle. **(E)** Initially unassigned density (unsharpened map) over the body domain of protein VIII, chain P (white frame). **(F)** Position and fit of the IIIa C-terminal residues identified by ModelAngelo into the E0 map density. Interaction of the C-terminal peptide of IIIa with VIII is possible in uncleaved VIII (E0, left), but after cleavage, protein VIII residues Thr64-Ala77 (highlighted in purple) partially block the binding site (E1, right).

We observed varying degrees of order in protein IIIa across the four structures compared, particularly affecting the protein N-terminus, the core-proximal domain, and the APD (**Figure 5A-C, Table S4**). The N-terminus (residues 2-29) was disordered exclusively in E0 particles (ANOVA p=1.11×10^-16^). Interestingly, this region is located close to the penton base N-terminus, which was also exclusively disordered in E0 (**Figure 3A and 5D**). The core-proximal domain tends to be more ordered when the core is present. Its long helix (Asp251-Phe275) is not ordered at all in E0, and the density for the turn starting at Phe275 is weaker in E1 than in F1. The APD can be traced both in E0 and F0, disconnected from the main body of the protein, but its density was weaker in E0. These observations indicate stabilization of the APD and core-proximal domain in the presence of the genome.

In E0 particles, an additional density strong enough to attempt interpretation (visible at 1.8σ) was found over the body domain of protein VIII, in the copy nearest to the center of the capsid facet (chain P), and disconnected from the rest of the proteins (**Figures 2 and 5E**). In F0 particles, it was tentatively assigned to part of the disordered N-terminal fragment of uncleaved VIII (12). Using ModelAngelo, we now determined that this small density corresponds to residues 478-486 of protein IIIa (**Figures 5B, 5F**), a peptide upstream of its AVP cleavage site. This identification reveals a previously unrecognized long-range interaction in which protein IIIa forms a bridge extending from the vertex to the facet center, much farther than previously appreciated. The pocket that accommodates the IIIa C-terminal region is partially occluded by residues Thr64–Ala77 of protein VIII following the latter’s extensive reorganization during maturation (**Figure 5F**), likely disrupting this interaction. Consistently, the density observed in E1 at a roughly corresponding location was of insufficient quality for tracing.

### Disordered contents in E0 particles

Although E0 and E1 particles lack genome and core proteins, they are not completely empty, as evidenced by the presence of weak, but higher than background, density inside the icosahedral shell (**Figure S3**). Previous studies on particles lacking the core attributed weak density accumulated near the inner capsid surface to the full-length (non-AVP processed) packaging protein L1 52/55 kDa (15). We attempted to find more detail about the organization of the weak density inside E0 particles, which contain unprocessed L1 52/55 kDa (**Figure S1**), by using less restrictive symmetry enforcement in our image processing (**Figure 6**). 3D alignment and classification of binned particles with no symmetry indicated variability in the amount and position of capsid contents (**Figure 6II**). We selected the class with the disordered contents more closely attached to the inner capsid surface for further processing (**Figure 6II, orange frame**), under the rationale that closer interactions with ordered components might facilitate order (and therefore observation) of some part of the protein generating the weak density. Since surface rendering suggested that density accumulation was stronger underneath the vertices (**Figure 6II, orange circle**), we used the orientations generated by the icosahedral refinement to extract and analyze subparticles corresponding to the vertex region, using localized reconstruction and classification. We did not observe any new density corresponding to ordered elements in any of the classes obtained, but one of them consisted of vertices lacking both pentons and peripentonal hexons (the whole GOS structure) (**Figure 6III, green circle**). We then reasoned that the lack of GOS might be related to some particular distribution of the disordered material inside the capsid. When particles that had contributed to the GOS-less class were binned and subjected to symmetry expansion and 3D classification without further alignment, we observed that the distribution of disordered material was asymmetrical in some classes, forming a shell thicker on one side of the capsid lumen, and again with a slight accumulation of material beneath the vertices (**Figure 6IV**). Failure to detect any ordered density that could shed light on the organization of full-length L1 52/55 kDa is consistent with the intrinsic disorder properties of this protein (27). The asymmetric distribution of disordered L1 52/55 kDa molecules within incomplete (GOS-less) E0 capsids is consistent with a concerted assembly pathway (**Figure 1**). In such a model, capsid assembly proceeds with directionality, with capsid fragments recruited sequentially around the core. Consequently, fragments incorporated early in the assembly process may have a different composition than those incorporated later, giving rise to the observed asymmetry. In any case, the subtle accumulation of disordered material, together with disorder in the N-terminal regions of protein IIIa and penton base, constitute three distinctive features uniquely observed in E0 particles, all of which converge in the region beneath the capsid vertex.

**Figure 6.**
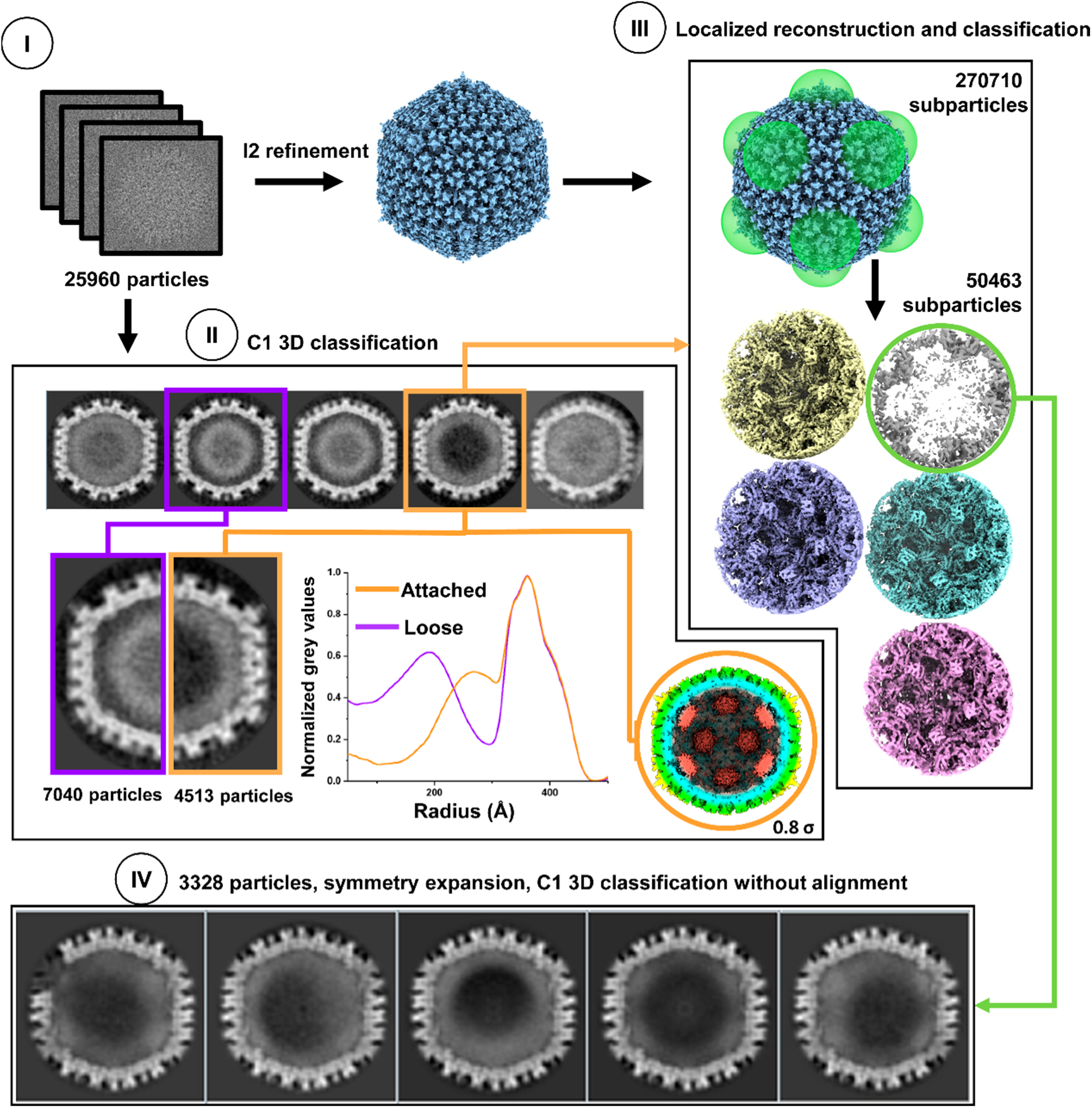
Internal disordered density in E0 particles. (I) Set of particles used for obtaining the E0 high-resolution map. (II) 3D alignment and classification of E0 particles without symmetry (binned to 5.4 Å/px). Frames and lines highlight classes corresponding to particles with disordered contents near the center of the capsid (purple) or attached to the inner side of the capsid (orange). Surface rendering colored by radius. (III) Localized reconstruction and classification of vertex subparticles extracted from the orange class in (II). The maps are viewed from inside the vertex. (IV) Examples of classes obtained from 3D classification without alignment (using icosahedral orientations and symmetry expansion) of particles contributing to the vertex class lacking GOS (green in (III), binned to 15.9 Å/px).

## Discussion

Unlike other extensively studied dsDNA viruses such as tailed phages or herpesvirus (28), adenoviruses do not undergo extensive capsomer rearrangements during morphogenesis. Our comparison between particles with and without genome shows only minimal structural changes related to the presence of dsDNA and accompanying core proteins. Only the core-proximal domain of protein IIIa is less ordered in E0 and E1 than in their counterparts F0 and F1, hinting at mobility in the absence of genome (**Figure 5A, B**). This observation differs from the results recently reported for HAdV-B7 VLPs, where in the absence of genome and core proteins both the N-terminal region of IIIa and most of protein VIII were largely disordered (29). It is currently unclear whether this is a trait specific to HAdV species B, or if it is caused by lack of some unidentified factor present in the natural infection but absent in the VLP recombinant production system. Conversely, changes related to proteolytic processing of packaging protein L1 52/55 kDa and internal minor coat proteins are notable, particularly affecting the connection of GOS elements to the rest of the capsid.

In the concerted assembly and packaging model (**Figure 1**), E0 particles represent failed assembly products where morphogenesis stopped in its early stages, before AVP, sliding on the dsDNA molecule, had time to start processing its targets (9, 15, 16). Residues 2-29 at the N-terminus of protein IIIa appeared disordered in E0 particles but well-ordered in the F0 and F1 genome-containing maps (**Figure 5A, B**). This behavior is not caused by the absence of genome, since the N-terminus is ordered in E1 particles, which are failed assembly products lacking the genome but having undergone some degree of processing by AVP. Additionally, the N-terminus of penton base, which is close to that of IIIa, appeared disordered exclusively in E0 particles (**Figure 3A, B**). Another characteristic feature of E0 particles is the presence of full-length, unprocessed L1 52/55 kDa protein forming a disordered shell that tends to accumulate under the vertices (**Figure 6**) (15), close to the region where the N-termini of penton base and IIIa appear disordered. Since neither penton base nor the N-terminus of IIIa are cleaved by AVP, but L1 52/55 kDa is (7), we propose that interactions with unprocessed L1 52/55 kDa are responsible for the disorder of these domains. That is, the presence of full-length L1 52/55 kDa would disorder protein regions cementing the icosahedral vertex in early assembly states (represented by E0). Disorder induced by L1 52/55 kDa in the penton base and IIIa N-terminal regions may play a role in weakening interactions at the vertex, to help correct assembly errors. As maturation proceeds, L1 52/55 kDa is cleaved by AVP and leaves the nascent capsid (7), allowing the N-termini of IIIa and penton base to become ordered (**Figure 7A**). The importance of weak protein interactions to avoid kinetic traps in the early steps of capsid assembly has been described for other dsDNA viruses, such as tailed bacteriophages (30).

**Figure 7.**
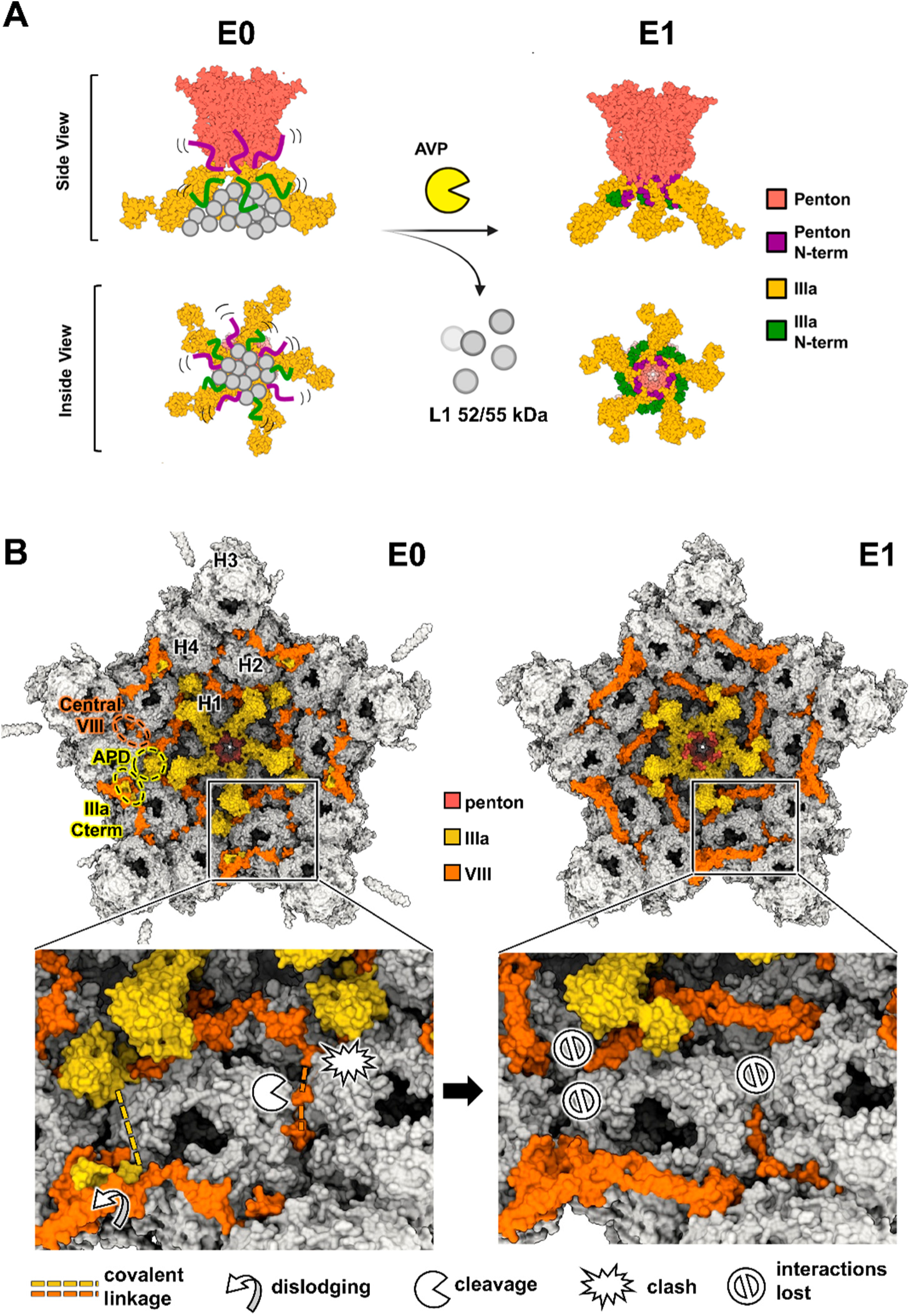
Structural changes underneath the adenovirus vertex upon maturation. (**A**) Cartoon illustrating how the N-termini in penton base and IIIa become ordered when L1 52-55 kDa is cleaved. **(B)** Connections between the GOS and the rest of the facet, established by the APD and the uncleaved proteins IIIa and VIII (E0, left), are disrupted upon AVP cleavage (E1, right).

Whereas the presence of uncleaved L1 52/55 kDa seems to induce disorder directly beneath the penton, changes in minor coat proteins IIIa and VIII affect the connections of the GOS with the rest of the capsid. The APD (residues 310-397) of protein IIIa appears ordered in both E0 and F0 particles (12), but not in E1 or F1 (3), indicating that its organization depends on maturation in HAdV-C5 (**Figure 5A-C**). Since the AVP cleavage site in protein IIIa is at residue 570 (9), maturation should not affect the appearance of the APD. In E0 and F0, the APD reinforces interactions at the GOS rim (**Figure 7B**). Additionally, we show that the uncleaved central peptide of protein VIII (residues 101-113) sits between hexons 2 and 4, reinforcing the second ring of capsomers around the vertex while also being covalently attached to the main body of VIII, since AVP processing has not occurred (**Figure 4 and S5**). Finally, a key finding of this work is the identification of a long-range connection established by protein IIIa that had been missed in previous studies. We identify residues 478-490 of protein IIIa interacting with the copy of protein VIII located beneath the GON (chain P, **Figure 5**). This identification implies that the C-terminal region of IIIa does not directly plunge into the core beneath the vertex as previously proposed (3), but instead, protein IIIa connects the vertex to the center of the facet *via* interactions with the distal copy of VIII. That is, before maturation, protein IIIa interacts with both copies of protein VIII in the AU, not just with the peripentonal one as previously observed.

These three stabilizing elements (APD, covalent bonding between the VIII central peptide and the VIII main body, and vertex-GON connection by the C-terminal region of IIIa) help connect proteins from the facet and the vertex when the virus particle has not been processed by AVP, but are remodeled during maturation (**Figure 7B**). AVP cleavage of protein VIII induces a movement of its head domain, causing it to overlap with the position of the APD in E0 and F0 particles. This conformational change in protein VIII causes clashes, displacing the APD from its position and making it non-traceable in E1 and F1 maps. This displacement doesn’t necessarily disorder the APD but blocks its interaction with hexons at the GOS rim. Concomitantly, the central region of VIII is separated from the main body of the protein, and residues 101-113 leave their attachment at the interface of hexons H2, H3, and H4, where they are replaced by the second excised peptide (residues 132-156), no longer covalently bound to the rest of the protein. The conformational change in protein VIII also displaces residues 478-490 of IIIa from their pocket in the copy of VIII closest to the facet center, disrupting the vertex-GON link established by this region in the immature capsid (**Figure 7B**).

Our findings reveal a more complex role of proteins IIIa and VIII in adenovirus stability modulation than previously realized. Protein VIII acts as a coiled spring, helping to keep capsid elements together during assembly while contributing to the removal of stabilizing interactions during maturation. In doing so, it facilitates the transition from a soft, stable particle to a brittle, metastable one, ready to start the uncoating cascade upon entry into the host cell (11, 31–33). Previous observations indicated that conformational plasticity of minor coat proteins is an important trait of adenovirus evolution (34, 35) as well as maturation (12). Here, we show how protein plasticity contributes to regulating vertex stability during morphogenesis, both to facilitate correct assembly and to prime the capsid for uncoating later in the infection cycle. Collectively, these findings position proteins IIIa and VIII as adenoviral counterparts of the scaffolding proteins that direct bacteriophage procapsid assembly (36), transiently organizing the interactions required for capsid construction before being remodeled or removed during maturation.

## Material and Methods

### Adenovirus production and purification

#### E0 particles

As a structural model for genomeless HAdV-C capsids lacking all AVP cleavages, HAdV-C2 *ts1* light particles were produced (37). A549 cells (ATCC: CCL-185) were grown in Dulbecco’s modified Eagle’s medium (DMEM) supplemented with 10% fetal bovine serum (FBS) (Sigma), 4 mM L-Glutamine (Sigma) and 1X non-essential amino acid solution (MEM Non-essential Amino Acid Solution, Sigma). To prevent contamination, 10 units-10 µg/mL penicillin-streptomycin and 0.05 mg/mL gentamicin were also added to the medium. Cells were seeded on p100 culture Petri dishes (Tissue culture-treated, Falcon) and maintained at 37⁰C in a humidified incubator with 5% CO_2_ to a density of 0.7-1 x10^6^ cells/mL (cell monolayer of 70% confluency). A seed consisting of 30 µL HAdV-C2 *ts1* (5.2 x 10^8^ viral particles (vp)/mL) was used for amplification in an increasing number of dishes at 32⁰C to produce infectious particles. The final infection to produce the immature particles was performed in 15 multilayer flasks (T500 multi-layer flask TripleFlask, ThermoFisher), at 39.5⁰C for 48 hours. Cells were centrifuged in a Megafuge 1.0 R centrifuge (Heraeus) for 40 minutes at 3345 x g at 4⁰C. Pellets were resuspended in 42 mL of medium from the infection supernatant and lysed by four freeze-thaw cycles. The remaining supernatant was kept for further virus purification.

For purification of intracellular virus, the cell lysate was centrifuged for 30 minutes at 3345 x g at 4⁰C to remove cellular debris. Viral particles were then separated by ultracentrifugation. The supernatant was distributed over six tubes (7 mL per tube) containing a discontinuous gradient of CsCl (1.4 g/mL and 1.25 g/mL in 2.5 mL phases) in TD 1X buffer (137 mM NaCl, 5.1 mM KCl, 0.7 mM Na_2_HPO_4_ x 7 H_2_O, 25 mM Tris-HCl pH 7.4). The gradients were then centrifuged in a Sorvall WX Ultra 100 (Thermo Scientific), using a SW41 Ti swinging bucket, at 240500 x g for 90 minutes at 18⁰C. After centrifugation, high– and low-density bands of virus particles were collected separately (approximately 1 mL per tube and band). High– or low-density pooled material (6 mL each) was loaded over 6 mL of 1.31 g/mL CsCl and ultracentrifuged for 18 hours at 240500 x g and 18⁰C. Bands collected from centrifugation were either dialyzed against HBS (20 mM HEPES pH 7.8, 0.15 M NaCl) using a semipermeable membrane (VSWP 0.025 µm filter, Millipore) (for volumes smaller than 1 mL) or transferred to Econo-Pac 10DG (Bio-Rad) disposable desalting columns for buffer exchange to HBS (for volumes larger than 1 mL). Virus concentration of aliquots collected from the column was estimated by absorbance (NanoDrop 1000 Spectrophotometer, Thermo Scientific) (measured at 280 and 260 nm). Aliquots were grouped by similar concentration and stored at –80 ⁰C with a final concentration of 10% glycerol. E0 capsids were collected from the lower density band, while the high-density band contained F0 particles.

For virus purification from the supernatant, 150 g of PEG8000 and 87.66 g of NaCl were added to each 1.5 L of DMEM and left stirring overnight at 4⁰C. The supernatant was then centrifuged for 20 minutes at 10,000 g and 4⁰C, and the pellets were resuspended in 10 mL of cold HBS. The collected solution was distributed into 2 tubes (7 mL per tube) containing a discontinuous CsCl gradient (1.4 g/mL and 1.25 g/mL in 2.5 mL phases) in TD 1X buffer (137 mM NaCl, 5.1 mM KCl, 0.7 mM Na_2_HPO_4_ x 7 H_2_O, 25 mM Tris-HCl pH 7.4). The rest of the procedure and storage were carried out as for intracellular virus purification.

#### E1 particles

As a structural model for genomeless HAdV-C capsids that have undergone partial maturation by AVP, HAdV-C5 FC31 light particles were produced. These light particles were referred to as L3 particles in (15). The HAdV-C5 FC31 RFP construct (19), has deletions in the E1 and E3 transcription units but is wild type for the proteins forming the capsid. Virus production and purification were performed as previously reported (15). HEK293 cells (ATCC: CRL-1573) were grown as described above for A549 cells. Once the cell monolayer reached approximately 70% confluence, purified virus was mixed with fresh medium (with 2% instead of 10% FBS) and added to the cells at a multiplicity of infection (MOI) of 5. Infected cells from 80 p100 Petri dishes were collected after 56 hours post-infection (hpi) and centrifuged for 40 min at 3345 x g at 4⁰C. Cell pellets were resuspended in 42 mL of medium from the infection supernatant and lysed by four freeze-thaw cycles. Intracellular virus was purified and stored as described for HAdV-C2 *ts1* particles.

#### Virus particle quantification and characterization

Virus particle concentration was quantified using the hexon protein emission spectra measured with a Fluorescence Spectrophotometer F-7000 (Hitachi). Samples of 150 µL in quartz cuvettes were excited at 285 nm, and emission was measured from 310 to 375 nm (emission maximum at 333 nm). Concentration was determined by calculating a calibration curve with a viral sample of known concentration.

For protein composition analysis via protein electrophoresis, virus samples were denatured in loading buffer (1% SDS, 1% β-mercaptoethanol, 10% glycerol, 50 mM Tris-HCl pH 6.8, 1.6% bromophenol blue) and incubated at 99⁰C for 15 minutes. 12% acrylamide gels or precast 4-20 % gradient acrylamide gels were run in a minigel Bio-Rad electrophoresis system. DualColor (Bio-Rad) was used as a molecular weight marker. Electrophoresis was carried out for approximately 1.5 hours (until the marker bands were easily distinguishable) at 100V in TBE buffer (89 mM Tris base, 89 mM boric acid, and 2 mM EDTA pH 8.5). After electrophoresis, silver staining was used to observe the protein bands. Gels were fixed in 20 mL ethanol, 5 mL acetic acid, and milli-Q water up to 50 mL for 30 minutes. Then, gels were sensitized in a solution of 15 mL ethanol, 5 mL sodium thiosulfate (5% w/v), 3.4 g of sodium acetate, and milli-Q water up to 50 mL for 30 minutes. After three washes of 3 minutes in milli-Q water, gels were stained in a 0.25% silver nitrate solution for 20 minutes, then washed again twice in milli-Q water for 30 seconds. Finally, to reveal the staining, gels were soaked in 50 mL milli-Q water with 1.25 g Na_2_CO_3_ and 20 µL formaldehyde 37%, for 5-10 minutes, until protein bands could be distinguished. To stop the staining process, gels were immersed in a 40 mM EDTA-Na_2_x2H_2_O solution. Approximately 6.5×10^9^ vp were loaded in each well of the gel for silver staining.

For Western blot analyses to determine the maturation state of proteins L1 52/55 kDa and VI, proteins were transferred from unstained gels to a nitrocellulose membrane using a Bio-Rad semi-dry transfer device (Trans-Blot Semi-dry) for 20 minutes at 15 V. Membranes were blocked with 5% powdered milk in Tris-buffered saline (TBS) (0.09% NaCl, 0.01% Tween 20, 0.1 M Tris-HCl pH 7.5) for 1 hour at room temperature and incubated overnight at 4 ⁰C with the primary antibody diluted in TBS with 0.5% milk powder. Afterwards, membranes were washed three times for 10 minutes in TBS and incubated at room temperature for 1 hour with the secondary antibody conjugated with horseradish peroxidase. Bands were revealed using the LiteAblot kit (Euroclone). Approximately 9.7×10^9^ vp were loaded in each well of the gel for Western blot. Primary antibodies were: rabbit anti-L1 52/55 kDa serum (1:2000 dilution) (38), and rabbit anti-VI serum (1:500 dilution) (39). Antibodies were kindly provided by Professors Patrick Hearing (Stony Brook University) and Urs Greber (Institute of Molecular Life Sciences-University of Zurich), respectively. As secondary antibody we used Peroxidase-conjugated Goat Anti-Rabbit (Jackson ImmunoResearch Laboratories) (1:100000 dilution).

### Cryo-EM sample preparation and image acquisition

Purified E0 particles were dialyzed into PBS (137 mM NaCl, 2.7 mM KCl, 8 mM Na_2_HPO_4_, and 2 mM KH_2_PO_4_, pH 7.8) for 1 hour at 4 ⁰C, and concentrated by centrifugation (10 minutes at 4 ⁰C) in an Amicon Ultra-0.5 Centrifugal Filter Unit (100 kDa, Millipore), for a final concentration of 2.1×10^13^ vp/mL. Concentrated virus particles were incubated on glow-discharged Quantifoil R 2/2 Copper/Rhodium (Cu/Rh) grids. Six consecutive incubations of the grids with 3 µL drops of sample followed by blotting were performed as described (40) to improve the number of particles lying in the carbon holes. Grids were vitrified in a Vitrobot (FEI) device. Purified E1 particles were dialyzed and concentrated similarly to E0 capsids, with a final concentration of 1.2×10^13^ vp/mL. Grids were prepared as for E0 particles, except that a Leica CPC device was used for vitrification. Grids were examined in a 200 kV Talos Arctica (FEI) microscope at the CNB-CSIC cryo-EM facility to assess particle concentration, ice quality, and overall sample suitability. Data acquisition for E0 particles was carried out at the European Synchrotron facility (ESRF, Grenoble). For the E1 sample, data was acquired at eBIC (Electron Bio-Imaging Centre, Diamond Light Source, Oxford) (**Table S1**).

### Image processing

Cryo-EM data processing was performed using the Scipion framework (41). After motion correction with Motioncor2 (42), the contrast transfer function (CTF) was estimated using CTFFIND4 (43), and low-resolution micrographs were discarded. Particles were picked using Xmipp (44) and extracted into 952-pixel boxes. Particles were downsampled by a factor of 2 to reduce computational resource usage before performing 2D and 3D classification processes with RELION (45), applying icosahedral symmetry. After 2D classification, low-quality particles were removed, and the rest were 3D-classified into three classes with an initial angular sampling of 3.7 degrees, which was gradually decreased to 0.5 degrees. The mature HAdV-C5 cryo-EM map (20) was used as an initial reference, filtered to 60 Å resolution. The largest class, also yielding the highest resolution, was further refined using RELION auto-refine with the initial 952-pixel boxes, followed by CTF refinement (beam tilt, anisotropic magnification, and per particle defocus) (46) and correction for the Ewald sphere curvature (47). The final resolution was estimated according to the gold-standard FSC= 0.143 criterion (48). The actual pixel size was calculated by fitting the HAdV-C5 asymmetric unit model (20) into the map in UCSF Chimera (49, 50). Map radial averages were calculated with Xmipp, using unsharpened maps filtered down to 6 Å resolution.

### Quantification of grey values

To compare the grey values in different zones, one AU of the HAdV-C5 model (20) was fitted into the cryo-EM maps to be compared, after overlapping them with UCSF Chimera (49). Markers were then placed over different domains of the adenovirus capsid proteins following the molecular model as a guide, and map values were measured at marker positions for all our experimental and previously published maps (E0, E1, F0, and F1). The same set of markers was used to measure grey values in all the maps to be compared. Grey values for each map were normalized to the median of the values obtained for hexon, the protein showing the highest intensity in all the maps. Normalization of the data, graph elaboration, statistical analyses, and hypothesis tests (ANOVA and Tukey mean difference) were conducted using Origin 2018 (https://www.originlab.com/).

### Non-icosahedral image processing

Particles used for obtaining the high-resolution E0 map were downsampled (binning 4) and 3D classified into five classes with C1 symmetry (7.5 degrees of angular sampling) using RELION (45), and separated according to the position of weak densities inside the capsid. Particles assigned to classes with a weak density shell closest to the inner capsid surface were selected and used for localized reconstruction of the vertex region as follows: the orientations determined for high-resolution icosahedral refinement were used to localize and extract vertices in 200 px box subparticles using LocalRec (51). Subparticles were 3D classified into five classes without further alignment in C1 symmetry. Virus particles with vertices assigned to the class lacking GOS were downsampled (binning 10) and subjected to icosahedral symmetry expansion to create 60 symmetry mates for each one of them using RELION. Finally, a 3D classification into 12 groups with C1 symmetry without performing particle alignment was carried out.

### Map interpretation

The icosahedral asymmetric units of E0 and E1 particles were built starting from the HAdV-C5 mature virus structure (PDB: 6b1t) (20). In E0 particles, the hexon and penton molecules were replaced by those experimentally determined for HAdV-C2 (PDB: 1p2z (52) and PDB: 1×9p (53)). Each asymmetric unit was then fitted to the map using UCSF Chimera (49), and iteratively refined using Coot (54) and Phenix real space refine (55), all within Scipion (56). Sequence alignments between HAdV-C2 and –C5 were performed using the protein BLAST alignment tool from NCBI (57) and the Geneious aligner included in Geneious Prime (https://www.geneious.com/). Sequence variations between HAdV-C2 and –C5 in the minor coat proteins in the asymmetric unit were introduced manually in Coot.

To analyze unassigned densities, remnant maps were constructed by masking the initial cryo-EM map with the molecular model of the residues already traced. UCSF Chimera, ChimeraX (50) and Coot were used for visualization. ModelAngelo (24) was applied to either the original sharpened or the remnant maps to identify the central peptide of VIII and the C-terminal region of IIIa. Initially, we provided no sequence to ModelAngelo and aligned the peptide assigned to the density with the HAdV-C2/5 genomes (NCBI Reference Sequences: NC_001405.1 and AC_000008.1, for HAdV-C2 and –C5 respectively) using BLAST (https://blast.ncbi.nlm.nih.gov). Then, ModelAngelo was run again, providing the sequence of the full-length matching proteins as starting point.

Root-mean-square deviation (RMSD) values for comparing proteins from different specimens were calculated using UCSF Chimera, considering all atoms after aligning the structures with the *matchmaking* tool.

## Data availability

The E0 and E1 maps and models have been deposited at the Protein Data Bank (PDB, https://www.ebi.ac.uk/pdbe/) and the Electron Microscopy Data Bank (EMDB, http://www.ebi.ac.uk/pdbe/emdb) with accession numbers 9TWZ / EMD-56372 (E0) and 9TVV / EMD-56354 (E1).

## Acknowledgements

Work supported by grants from the Spanish State Research Agency, with co-funding from the European Regional Development Fund (BFU2016-74868-P/AEI/10.13039/501100011033, PID2019-104098GB-I00/AEI/10.13039/501100011033 and PID2022-136456NB-I00/AEI/10.13039/501100011033). The C.S.M. group is a member of the Spanish Adenovirus Network (RED2022-134221-T/AEI/10.13039/501100011033). The CNB-CSIC was further supported by AEI Severo Ochoa Excellence grants SEV-2017-0712/AEI/10.13039/501100011033 and CEX2023-001386-S/AEI/10.13039/501100011033). J. G. held a predoctoral contract (BES-2017-079868/10.13039/501100011033) funded by MICIU/AEI and by ESF Investing in your future.

We acknowledge excellent technical support by the CNB-CSIC Electron and Cryo-electron Microscopy facilities, as well as Titan Krios data collection with Katie Cunnea at Diamond Light Source UK national electron Bio-Imaging Center (eBIC) under proposal EM15997-28, and Gregoy Effantin at the European Synchrotron Radiation Facility (ESRF) under proposal MX-2263 (https://doi.org/10.15151/ESRF-ES-267184482).

## Author contributions

C.S.M. designed research. J.G. carried out most of the experimental research and data analysis, with help from M.M. and R.M. J.G. and C.S.M. wrote the paper, with input from all other authors.

