## Supplementary Figures and Tables for "Protease-driven remodeling of stabilization networks during adenovirus assembly"

[2] Escuela Politécnica Superior. Universidad Autónoma de Madrid. Madrid (Spain)

**Short title: Structural remodeling during adenovirus assembly**

**This PDF file includes:**

Supplementary figures S1 to S5

Supplementary tables S1 to S4

Supplementary references

### Supplementary Figures

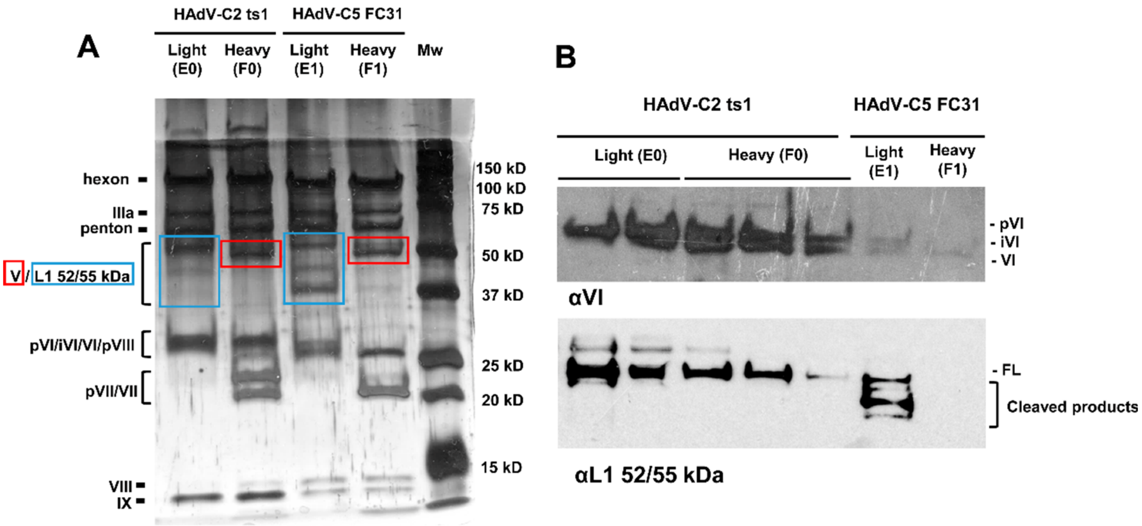


**Figure S1. Protein composition of E0 and E1 particles and their genome-containing counterparts**. Heavy, mature particles produced by HAdV-C5 FC31 are used here for F1. (A) Silver-stained SDS-PAGE. Mw: molecular weight ladder. (B) Western blot using specific antibodies to proteins VI and L1 52/55 kDa. Prefix "p" indicates precursor versions of capsid and core AVP targets. iVI: intermediate form cleaved at one of the two AVP sites. FL: full-length L1 52/55 kDa.

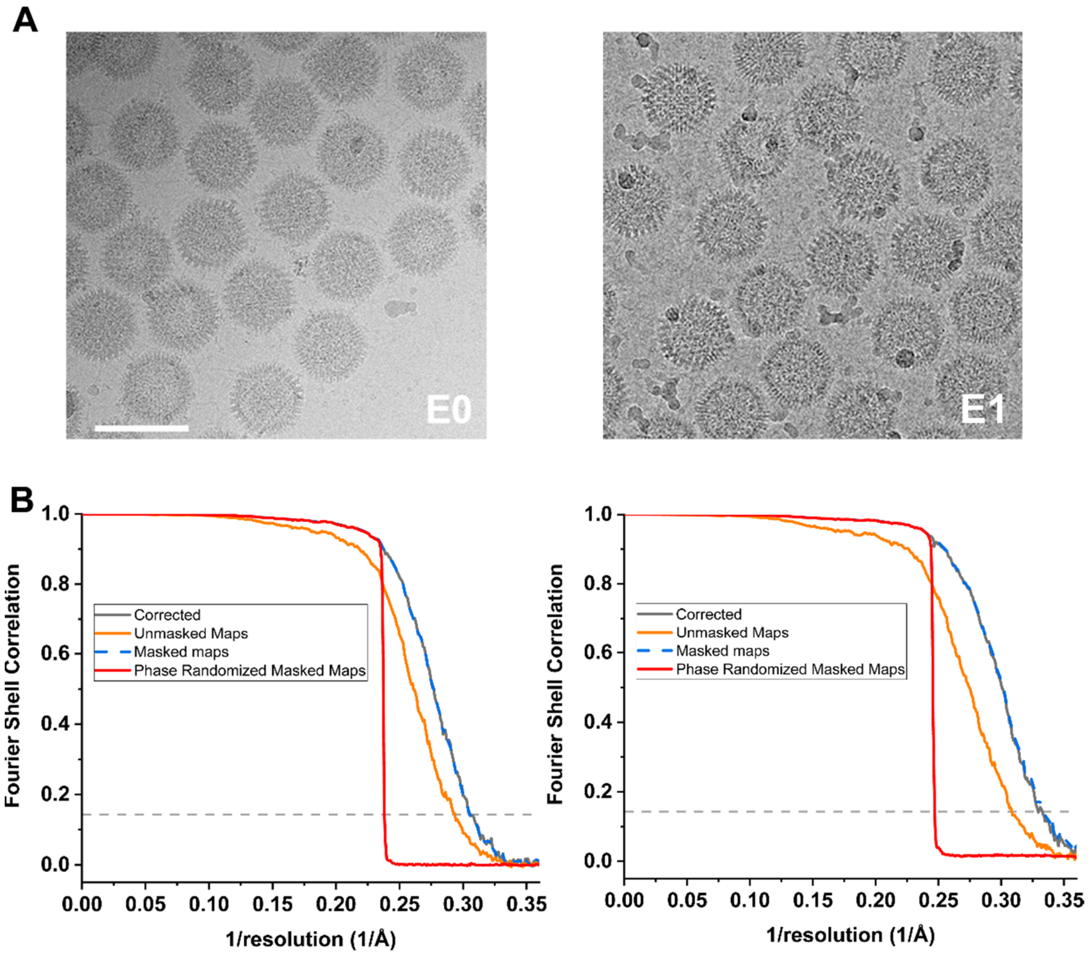


**Figure S2. Cryo-EM single-particle averaging of E0 and E1 particles.** (A) Example micrographs. Scale bar: 100 nm. (B) Fourier Shell Correlation (FSC) curves showing final resolutions of 3.3 (E0, left) and 3.0 Å (E1, right). Note that the corrected FSC curves overlap with those for the masked maps.

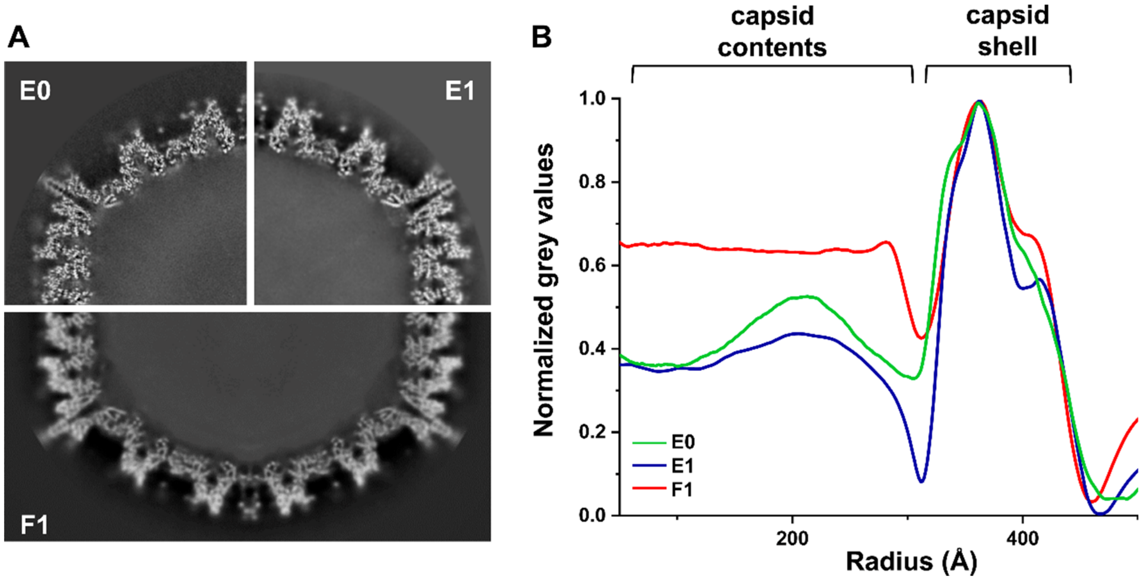


**Figure S3. Capsid contents of the E0, E1, and F1 maps.** (A) Central slice of cryo-EM maps of E0, E1, and F1 particles. A HAdV-C5 map from (Hernando-Pérez, Martín-González et al., 2020) was used here as F1 because it retains the core density, unlike the F1 map from (Dai, Wu et al., 2017) where the core was masked during refinement. (B) Normalized radial average profiles of the cryo-EM maps shown in (A).

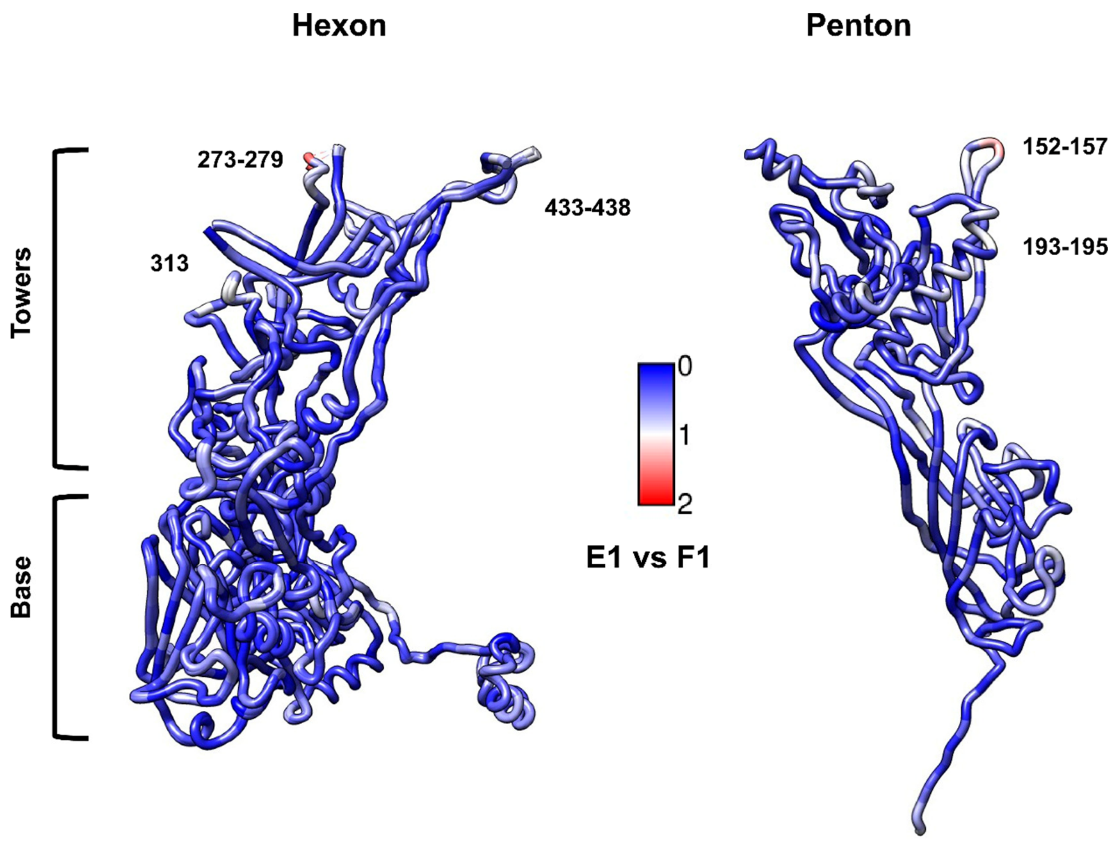


**Figure S4. Comparison between hexon and penton base monomers in E1 and F1 particles.** Scale bars indicate RMSD in Å. Residue numbers indicate the regions with the largest differences between specimens.

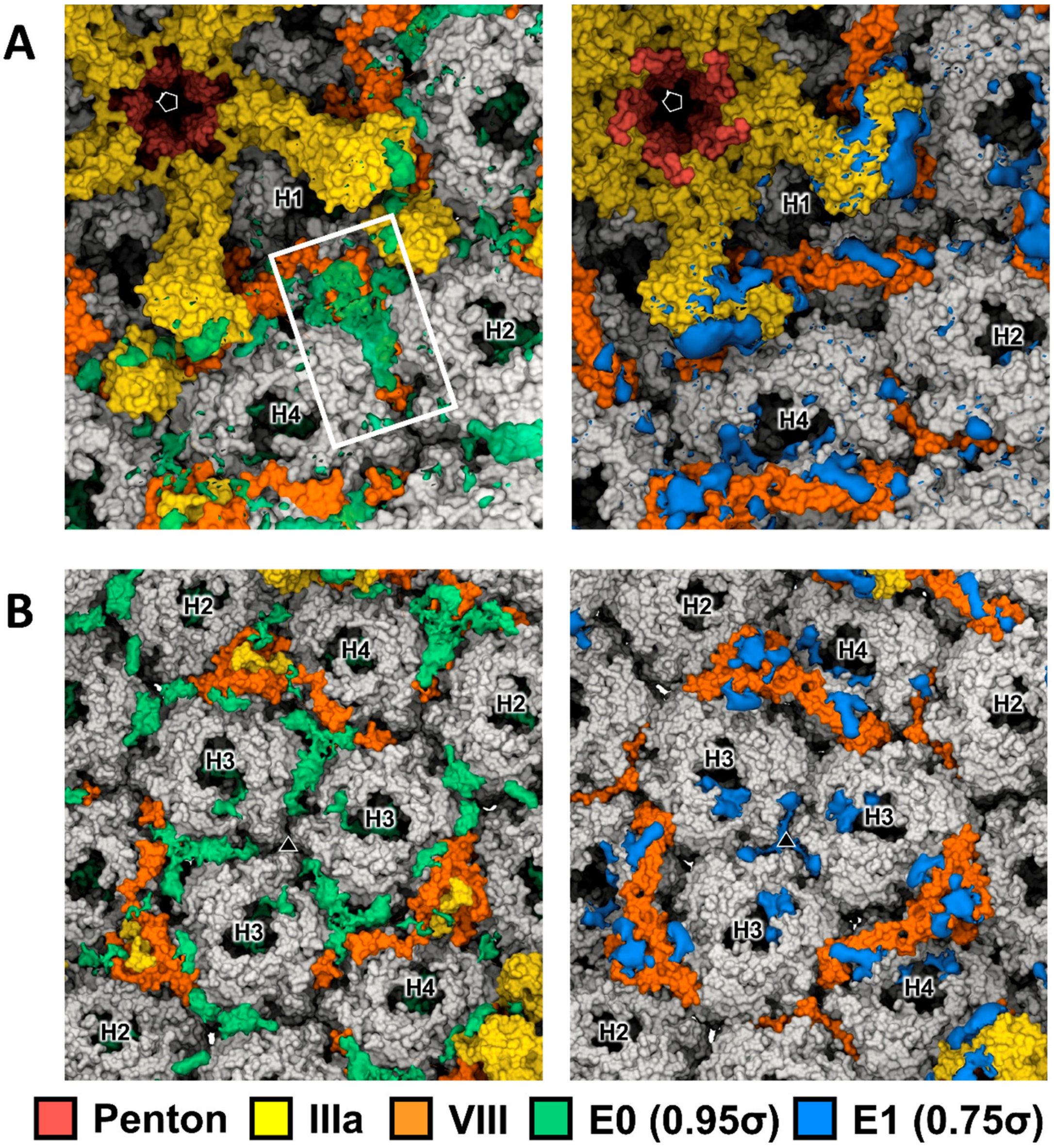


**Figure S5. Remnant weak densities near protein VIII.** (A) Weak density (white frame) connecting the copy of protein VIII beneath the GOS (chain O) and the central peptide in E0 (left). No connecting weak density is observed in E1 (right). (B) Densities near the protein VIII copy beneath the GON (chain P). Triangle: 3-fold icosahedral axis.

### Supplementary Tables

**Table S1.** Cryo-EM data collection and processing parameters.

| | **E0 particles** | **E1 particles** |
|---|---|---|
| Microscope | Titan Krios (ESRF) | Titan Krios (eBIC) |
| Voltage (kV) | 300 | 300 |
| Camera | Gatan K2 | Falcon III |
| Magnification | 105,000 | 95,890 |
| Nominal pixel size (Å) | 1.35 | 1.46 |
| Final pixel size (Å) | 1.33 | 1.38 |
| Electron dose (e-/Å$^2$) | 21.7 | 38.6 |
| Dose/frame (e-/Å$^2$) | 0.90 | 0.99 |
| Frames per movie | 24 | 39 |
| Defocus range (µm) | -1.5 to -3.0 | -1.5 to -3.0 |
| Micrographs | 8398 | 4829 |
| Initial particles | 33206 | 39125 |
| Final particles | 25960 | 30228 |
| Symmetry | Icosahedral | Icosahedral |
| B-factor (Å$^2$) | -127.83 | -179.73 |
| Final resolution | 3.3 Å | 3.0 Å |

**Table S2.** Validation statistics for the models traced on the E0 and E1 maps.

| | E0 particles | E1 particles |
|---|---|---|
| **Composition** | | |
| Chains | 29 | 25 |
| Atoms | 98686 | 97893 |
| Residues (amino acids) | 12404 | 12239 |
| **Validation** | | |
| Bonds (RMSD); outliers > 4 σ | | |
| Length (Å) | 0.004; 0 | 0.007; 0 |
| Angles (˚) | 1.014; 10 | 1.061; 16 |
| Molprobity score | 2.09 | 2.22 |
| Clash score | 11.19 | 10.68 |
| Ramachandran plot (%) | | |
| Outliers | 0.02 | 0.01 |
| Allowed | 3.51 | 3.37 |
| Favored | 96.48 | 96.62 |
| Rotamer outliers (%) | 2.41 | 3.95 |
| **Model vs. Data** | | |
| CC (mask) | 0.84 | 0.85 |
| CC (box) | 0.42 | 0.50 |
| CC (volume) | 0.79 | 0.81 |
| CC (peaks) | 0.21 | 0.32 |
| CC (main chain) | 0.84 | 0.85 |
| CC (side chain) | 0.82 | 0.83 |

**Table S3 (next page).** Proteins traced in the E0 and E1 maps. Each chain is identified by three letters (XYZ), where X denotes the icosahedral asymmetric unit, Y is the standard chain identifier in PDB files (a single character, for example, O and P for the two copies of protein VIII), and Z is a single character protein identifier (for example, p for penton base, h for hexon, 3 for protein IIIa, 6 for protein VI, etc). NA: not available.

| | E0 particle | | | E1 particle | | |
|---|---|---|---|---|---|---|
| **Protein** | **Length (aa)** | **Chain ID** | **Residues traced** | **Length (aa)** | **Chain ID** | **Residues traced** |
| **Hexon** | 968 | 0Ah | 5-137, 172-194, 204-262, 270-280, 291-443, 454-965 | 952 | 0Ah | 4-138, 164-253, 258-273, 279-433, 437-950 |
| | | 0Bh | 6-138, 172-194, 204-263, 269-280, 291-443, 454-965 | | 0Bh | 6-138, 164-253, 258-273, 279-433, 437-950 |
| | | 0Ch | 7-137, 173-194, 204-262, 270-280, 291-443, 454-963 | | 0Ch | 1-138, 164-253, 258-273, 279-433, 437-947 |
| | | 0Dh | 6-138, 171-193, 203-262, 269-280, 291-443, 454-966, | | 0Dh | 2-138, 164-253, 258-273, 279-433, 437-952 |
| | | 0Eh | 5-139, 171-194, 203-262, 270-280, 291-443, 454-964 | | 0Eh | 7-138, 164-253, 258-273, 279-433, 437-949 |
| | | 0Fh | 6-138, 171-194, 203-262, 269-280, 291-443, 454-966 | | 0Fh | 6-138, 164-253, 258-273, 279-433, 437-952 |
| | | 0Gh | 6-137, 171-194, 204-262, 270-280, 291-443, 454-963 | | 0Gh | 6-138, 164-253, 258-273, 279-433, 437-952 |
| | | 0Hh | 3-138, 171-194, 203-262, 268-280, 291-443, 454-963 | | 0Hh | 1-138, 164-253, 258-273, 279-433, 437-950 |
| | | 0Ih | 6-138, 171-194, 203-262, 270-280, 291-443, 454-966 | | 0Ih | 6-138, 164-253, 258-273, 279-433, 437-951 |
| | | 0Jh | 6-138, 171-194, 204-262, 270-280, 291-443, 454-966 | | 0Jh | 5-138, 164-253, 258-273, 279-433, 437-951 |
| | | 0Kh | 6-138, 171-194, 204-262, 270-280, 291-443, 454-966 | | 0Kh | 2-138, 164-253, 258-273, 279-433, 437-951 |
| | | 0Lh | 3-138, 172-194, 204-262, 270-280, 291-443, 454-963 | | 0Lh | 2-138, 164-253, 258-273, 279-433, 437-950 |
| **Penton** | 571 | 0Mp | 51-296, 376-569 | 571 | 0Mp | 37-296, 377-571 |
| **IIIa** | 585 | 0N3 | 30-215, 222-273, 315-353, 366-392, 476-490 | 585 | 0N3 | 4-216, 226-277, 295-302 |
| **VIII** | 227 | 0O8 | 1-63, 101-113, 146-227 | 227 | 0O8 | 2-111, 132-156, 158-227 |
| | | 0P8 | 2-64, 143-227 | | 0P8 | 2-111, 158-227 |
| **IX** | 140 | 0Q9 | 7-57, 67-129 | 140 | 0Q9 | NA |
| | | 0R9 | 7-54, 94-133 | | 0R9 | NA |
| | | 0S9 | 7-57, 95-138 | | 0S9 | NA |
| | | 0T9 | 7-56, 64-82, 96-130 | | 0T9 | NA |
| **VI** | 250 | 0a6 | 5-33 | 250 | 0a6 | 6-33 |
| | | 0b6 | NA | | 0b6 | NA |
| | | 0c6 | 5-42 | | 0c6 | 5-29 |
| | | 0d6 | 5-33 | | 0d6 | 5-33 |
| | | 0e6 | 5-33 | | 0e6 | 5-33 |
| | | 0f6 | 5-13 | | 0f6 | 5-12 |
| | | 0g6 | 5-19 | | 0g6 | 5-29 |
| | | 0h6 | NA | | 0h6 | NA |
| | | 0i6 | 5-29 | | 0i6 | 5-28 |
| | | 0j6 | 5-32 | | 0j6 | 5-29 |
| | | 0k6 | NA | | 0k6 | NA |
| | | 0l6 | 5-31 | | 0l6 | 5-12 |

**Table S4.** Quantification of density differences in selected regions of the cryo-EM maps. Results of Tukey's Honestly Significant Difference (HSD) post-hoc test for pairwise comparisons between group means. *N* is the number of pixels in which the gray values were measured. *MeanDiff* represents the difference between the means of the two groups being compared. *SEM* is the standard error of the mean difference. The *q Value* corresponds to the studentized range statistic used to assess differences among multiple groups. *P* denotes the adjusted p-value for each comparison, and *Alpha* is the significance threshold. *Sig* indicates whether the difference is statistically significant (*True* if *P* < *Alpha*). *LCL* and *UCL* represent the lower and upper bounds of the confidence interval for the mean difference; intervals that do not include zero indicate statistically significant differences (shaded cells).

| Region (residues) | N | Maps compared | MeanDiff | SEM | q Value | P | Alpha | Sig | LCL | UCL |
|---|---|---|---|---|---|---|---|---|---|---|
| Penton base N-term (37-51) | 19 | E1 E0 | 0.70422 | 0.06991 | 14.24535 | 5.9517E-9 | 0.05 | True | 0.52035 | 0.88809 |
| | | F0 E0 | 0.53101 | 0.06991 | 10.74146 | 0 | 0.05 | True | 0.34713 | 0.71488 |
| | | F0 E1 | -0.17322 | 0.06991 | 3.50389 | 0.07209 | 0.05 | False | -0.35709 | 0.01066 |
| | | F1 E0 | 0.47867 | 0.06991 | 9.68287 | 0 | 0.05 | True | 0.2948 | 0.66255 |
| | | F1 E1 | -0.22555 | 0.06991 | 4.56248 | 0.00998 | 0.05 | True | -0.40942 | -0.04167 |
| | | F1 F0 | -0.05233 | 0.06991 | 1.05859 | 0.877 | 0.05 | False | -0.2362 | 0.13154 |
| IIIa N-term (2-29) | 15 | E1 E0 | 0.68281 | 0.05454 | 17.70642 | 0 | 0.05 | True | 0.5301 | 0.83552 |
| | | F0 E0 | 0.52587 | 0.05454 | 13.63682 | 0 | 0.05 | True | 0.37316 | 0.67858 |
| | | F0 E1 | -0.15693 | 0.05454 | 4.0696 | 0.04103 | 0.05 | True | -0.30964 | -0.00423 |
| | | F1 E0 | 0.59902 | 0.05454 | 15.5338 | 0 | 0.05 | True | 0.44632 | 0.75173 |
| | | F1 E1 | -0.08378 | 0.05454 | 2.17262 | 0.54281 | 0.05 | False | -0.23649 | 0.06893 |
| | | F1 F0 | 0.07315 | 0.05454 | 1.89698 | 0.66656 | 0.05 | False | -0.07956 | 0.22586 |
| IIIa APD (310-397) | 29 | E1 E0 | -0.30431 | 0.02044 | 21.05324 | 0 | 0.05 | True | -0.3608 | -0.24781 |
| | | F0 E0 | 0.38428 | 0.02044 | 26.58586 | 4.75247E-8 | 0.05 | True | 0.32778 | 0.44077 |
| | | F0 E1 | 0.68858 | 0.02044 | 47.6391 | 1.25812E-7 | 0.05 | True | 0.63209 | 0.74507 |
| | | F1 E0 | -0.29214 | 0.02044 | 20.21139 | 0 | 0.05 | True | -0.34863 | -0.23565 |
| | | F1 E1 | 0.01217 | 0.02044 | 0.84185 | 0.9756 | 0.05 | False | -0.04432 | 0.06866 |
| | | F1 F0 | -0.67641 | 0.02044 | 46.79725 | 1.23005E-7 | 0.05 | True | -0.73291 | -0.61992 |
